# Effects of stimulus modality and response type on oddball stimulus discrimination using polarity-considered EEG microstate labeling

**DOI:** 10.1101/2025.04.29.650929

**Authors:** Tatsumi Tsubaki, Shiho Kashihara, Tomohisa Asai, Hiroshi Imamizu, Isao Nambu

**Affiliations:** Graduate School of Engineering, Nagaoka University of Technology, 1603-1 Kamitomioka, Nagaoka City, Niigata 940-2188, Japan; Department of Cognitive Neuroscience, Cognitive Mechanisms Laboratories, Advanced Telecommunications Research Institute International (ATR), Keihanna Science City, Kyoto 619-0288, Japan; Department of Psychology, Graduate School of Humanities and Sociology and Faculty of Letters, The University of Tokyo, 7-3-1 Hongo, Bunkyo-ku, Tokyo 113-8654, Japan

**Author notes:** (Faculty of Humanities, Fukuoka University, 8-19-1 Nanakuma, Jonan-ku, Fukuoka City, Fukuoka 814-0180, Japan.) (Shiho Kashihara).

**Keywords:** EEG microstates, oddball task, classification

## Abstract

**Objective:** Brain–computer interfaces (BCIs) require effective feature extraction and dimensionality reduction from multidimensional brain signals. Electroencephalogram (EEG) microstate analysis offers a fast and noise-resistant approach by classifying the states of brain signals into spatial distribution patterns (templates). Each EEG segment was assigned the template with the highest spatial correlation, reducing the information to a one-dimensional representation. However, prior BCI studies have often ignored the polarity of spatial distributions in these templates. Incorporating polarity during labeling may enhance classification performance. This study investigated the effectiveness of polarity-considered microstate labeling for classifying infrequent stimuli in an auditory-visual oddball task with implications for BCI applications.

**Method:** EEG recordings were analyzed using polarity-considered microstate labeling to classify infrequent stimuli. This study examined the effects of stimulus modality (auditory or visual), modality conditions (unimodal: stimulus and response in the same modality; cross-modal: stimulus and response in different modalities), and response type (key-press task vs. mental counting task) on classification accuracy. Machine learning models were used for classification, including support vector machine, random forest, logistic regression, XGBoost, CatBoost and K-means methods.

**Results:** Polarity-considered labeling outperformed the non-polarity approach, especially in decision-tree-based models (20.1% improvement in the key-press task and 22.2% improvement in the mental counting task). A significant interaction was observed between stimulus modality and response type, with the highest accuracy achieved when the infrequent stimuli in the key-press task involved cross-modal visual information.

**Conclusion:** The findings suggest that polarity-considered microstate labeling enhances EEG-based classification. This approach has potential applications in BCI, such as in P300 spellers using cross-modal auditory-visual stimuli.

## 1. Introduction

Brain–computer interface (BCI) systems analyze brain signals to control external devices and hold promise for a wide range of applications, including medical care and daily life support. A key requirement for implementing BCI systems is the extraction and dimensionality reduction of features from high-dimensional brain signals. P300 speller is a character input system that utilizes event-related potentials (ERPs) elicited by focusing attention on flashing characters, as first proposed by Farwell and Donchin ^(1)^. Although BCI systems are considered practical for individuals with severe motor impairments, such as those with amyotrophic lateral sclerosis (ALS), they have not yet achieved widespread usability. A major barrier is the limited classification performance and detection accuracy of the P300 component in single-trial analyses. Farwell and Donchin ^(1)^, identified the Pz electrode (located over the parietal region) as the primary site for observing the P300 component. Similarly, Rezeika et al. ^(2)^ noted that “the P300 signal is usually intensified over the central parietal region of the brain and can be detected using electroencephalogram (EEG),” highlighting the reliance on localized brain activity. However, Krusienski et al. ^(3)^ demonstrated that using several electrodes improved classification accuracy. These findings suggest that expanding spatial coverage can enhance P300 detection performance.

EEG microstate analysis classifies multi-channel EEG signals into spatial patterns (templates) and extracts the characteristics of each template as features. This approach facilitates interpreting brain activity during task performance. Moreover, employing microstate analysis with many electrodes makes it possible to enhance the interpretability of the brain states involved in EEG-based classification. This study applies EEG microstate analysis in a BCI system to detect P300 components and extract features reflecting more comprehensive brain dynamics.

### 1.1 Related Work and Distinction of the Present Study

Existing BCI studies utilizing EEG microstate analysis ^(4)-(7)^ have typically generated four to five microstate templates and extracted low-dimensional features, such as mean duration, frequency of occurrence, time coverage, and transition probability. For instance, Cui et al. ^(4)^ examined inter-subject variability in motor imagery BCI performance by analyzing four types of EEG microstates using the above metrics. Xiong et al. ^(5)^ improved the precision of microstate analysis by employing global map dissimilarity (GMD)- driven density canopy clustering, achieving a classification accuracy of up to 98.41% on a BCI dataset. However, to our knowledge, no previous studies have applied EEG microstate labeling to BCI systems, where each EEG time point is assigned the template with the highest spatial correlation. This labeling approach enables extracting one-dimensional features that are computationally simple and robust against noise—beneficial to the acceleration and practical deployment of future BCI systems.

### 1.2. Polarity in EEG Microstates

Previous studies ^(4)-(7)^ have typically ignored the polarity (positive/negative) of EEG signals in microstate analysis. However, Kashihara et al. ^(8)^ proposed that incorporating polarity into microstate labeling enables smoother transitions and improves the detection of age-related changes in brain dynamics. Based on this insight, we hypothesized that incorporating polarity into microstate templates by adding polarity-inverted versions of existing templates would allow extracting more informative features. Mahini et al. ^(9)^ suggested that the topographical changes observed in the oddball response, particularly components such as N2 and P3, may reflect polarity-inverted isomorphic states. Therefore, this study introduces polarity-considered microstate labeling to achieve more effective feature extraction. The motivation for considering polarity lies in the interpretation presented by Kashihara et al. ^(8)^, which suggests that polarity reflects the direction of neural sources, the structural constraints of the brain, and the excitatory/inhibitory (E/I) balance. Hence, including polarity may allow microstates to capture neural activity with greater physiological significance.

### 1.3. Experimental Factors in the Oddball Paradigm

The oddball paradigm ^(10)^ is a widely used cognitive task to study P300 responses in BCI research. This paradigm randomly presents infrequent target stimuli among frequent standard stimuli. We explored stimulus classification within the oddball paradigm to develop a BCI system incorporating polarity-considered microstate labeling to achieve higher classification accuracy. We hypothesized that refining the task environment would lead to further improvements in performance. Kashihara et al. ^(11)^, used three experimental factors in the oddball paradigm.

i. The modality of infrequent stimuli (auditory or visual)
ii. Combinations of frequent and infrequent stimuli (unimodal or cross-modal)
iii. The type of response to infrequent stimuli (key response task or count task)

Existing studies are inconsistent regarding whether visual stimuli ^(12, 13)^ or auditory stimuli ^(14)^ elicit stronger responses. Research on stimulus combinations ^(15, 16)^ suggests that late components, including the P300, tend to be enhanced under cross-modal conditions, whereas early components remain unaffected. Regarding response type, some studies have reported enhancement of the P300 with key response tasks ^(17, 18)^, others with count tasks ^(19, 20)^, and others have reported no significant difference ^(21)^. Nevertheless, from the perspective of classification performance in oddball-based BCIs, cross-modal combinations of frequent and infrequent stimuli offer potential advantages. These experimental factors are known to influence ERP components, and evidence suggests that polarity- considered microstate labeling can capture similar brain responses ^(11)^. Accordingly, applying polarity-based microstate labeling to classification tasks under these varied experimental conditions may enable us to propose optimal task settings for this method.

Therefore, the present study focused on the following three objectives: (I) to evaluate the classification performance of P300 responses using microstate labeling, which differs from conventional approaches; (II) to investigate the potential improvement in performance achieved by utilizing polarity-considered microstate labeling, a recent advancement in the field; (III) to examine the correspondence between classification performance under different experimental conditions in the oddball paradigm and the application of these labeling methods. For objective (I), we evaluated whether the classification performance using one-dimensional microstate sequences exceeds chance levels. While linear classifiers, such as linear discriminant analysis (LDA) and step-wise linear discriminant analysis (SWLDA), have been widely adopted for P300 classification, with SWLDA achieving over 90% accuracy in previous studies ^(3)^, this study compares a broader set of models. For objective (II), we evaluated whether polarity-considered labeling performs better than conventional polarity-ignored labeling. Additionally, we explore the neuroscientific underpinnings of this approach from the perspective of the topographical dynamics of brain activity. For objective (III), we investigated whether known differences in ERP responses under various oddball conditions, such as stimulus modality, frequency ratio, and response type, were reflected in the classification performance using polarity-considered microstate labeling. Based on these objectives, we discuss the implications of our results for future applications in the development of BCI systems.

## 2. Method

### 2.1. Experiment Data

For classification, we used data from the oddball task conducted by Kashihara et al. ^(11)^. Two datasets were collected from 20 healthy participants in their 20s and 30s. (Their studies were approved by the Ethical Committee of ATR (approval numbers: 21-144 and 21-143). The experiment was designed to examine the three factors described in Section 1.4. Four stimulus conditions were used: auditory only (unimodal auditory: uniA), visual only (unimodal visual: uniV), high-frequency visual with low-frequency auditory (cross-modal auditory-target: croA), and high-frequency auditory with low-frequency visual (cross-modal visual-target: croV). The modality conditions were counterbalanced across the participants. Auditory stimuli consisted of pure tones at 1000 Hz for frequent stimuli and at 2000 Hz for infrequent stimuli. The visual stimuli consisted of circles as frequent stimuli and stars as infrequent stimuli. Two task groups were defined for responses to infrequent stimuli: the key response task (keyresTask), in which participants (all right-handed) were instructed to press the spacebar with their right index or middle finger as quickly and accurately as possible, and the counting task (countTask), in which participants silently counted the number of target stimuli. In the countTask, the assignment of stimulus frequencies was also counterbalanced across participants. KeyresTask group included 12 females and eight males (mean age = 30.7 years, SD = 6.9), and the countTask group included 10 females and 10 males (mean age = 25.0 years, SD = 5.3). EEG recordings were performed using an R-Net 32-channel system (Brain Products GmbH, Gilching, Germany) and a BrainAmp MR plus amplifier (Brain Products GmbH). The 32 electrode positions followed the international 10-10 system (Fp1, Fp2, Fz, F3, F4, F7, F8, F9, F10, FC1, FC2, FC5, FC6, Cz, C3, C4, T7, T8, CP1, CP2, CP5, CP6, Pz, P3, P4, P7, P8, P9, P10, Oz, O1, and O2). Each trial consisted of a 200 ms stimulus presentation and a 1000 ms interstimulus interval. 320 frequent and 80 infrequent stimuli were presented, with infrequent stimuli pseudo-randomized to avoid consecutive occurrences. In keyresTask, trials containing errors, either false alarms (FA: incorrect key press) or misses (no response), were excluded from the analysis. After artifact rejection, the minimum numbers of valid frequent and infrequent trials across participants were 312 and 68, respectively. Further details can be found in Kashihara et al. ^(11)^.

### 2.2. Preprocessing

A bandpass filter from 1 to 45 Hz was applied to the EEG signals acquired in the experiment, followed by Artifact Subspace Reconstruction (ASR) ^(22)^ to remove nonstationary noise and the influence of bad electrodes. Next, average re-referencing and Independent Component Analysis using Adaptive Mixture ICA (AMICA) ^(23)^ were performed. Dipole fitting and component classification were conducted using ICLabel ^(24)^, and the components not classified as brain-related were removed. The EEG data were then segmented into epochs ranging from 200 ms before to 1000 ms after the stimulus onset.

An overview of the post-epoch data processing pipeline is shown in Fig. 1. The epoched data were again filtered with a 1–45 Hz band-pass filter, and the channel-wise mean was computed within the specified time windows. The resulting spatial distribution was correlated with the templates provided by Kashihara et al. ^(8)^, and template matching was performed using a winner-take-all approach.

**Fig. 1.**
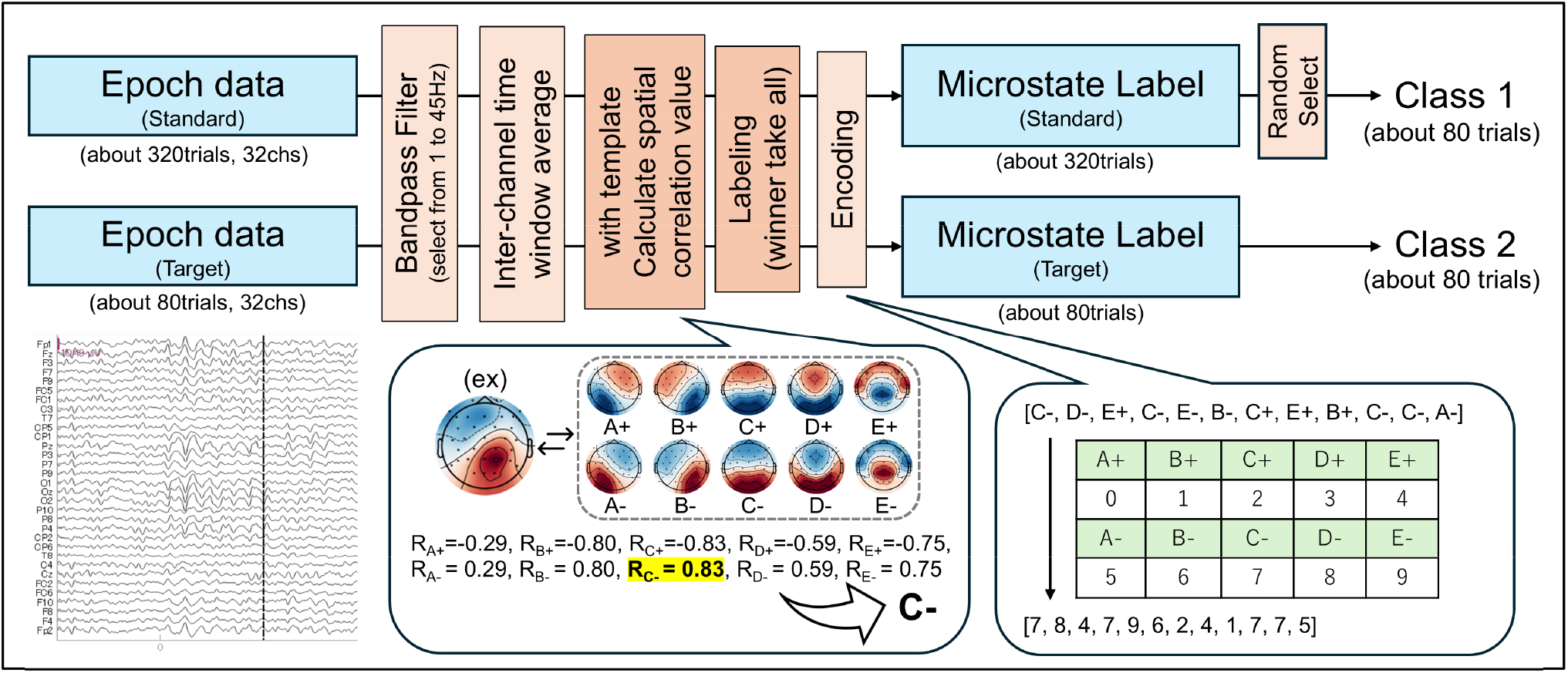
Decoding process. The process of microstate label coding from EEG epoch data, segmented from 200 ms before to 1000 ms after stimulus onset in a preprocessed oddball paradigm. The EEG topographies shown in the center represent a set of 10 templates derived from the LEMON dataset ^(25)^, obtained using modified k-means clustering with five clusters and their corresponding polarity-inverted maps.

The templates created by Kashihara et al. ^(8)^ were derived from the LEMON dataset ^(25)^ using modified k-means clustering, resulting in five spatial maps. By including polarity-inverted versions of each, a total of ten microstates (A± to E±) were obtained (Fig. 1, center). The template matching results were then encoded as integers from zero to nine. Finally, a random subset of frequent trials was selected to balance the dataset and equalize the number of trials between frequent and infrequent stimuli.

### 2.3. Label Occurrence Frequency

To investigate the temporal dynamics of the microstate labels obtained in Section 2.2, the occurrence frequency of each label is calculated within sliding time windows of 100 ms in size, with a 25% overlap. For each time window, the occurrence frequency was defined as the total count of each of the 10 template labels across all trials, divided by the total number of trials.

### 2.4. Classification

#### 2.4.1. Temporal generalization matrix

To evaluate temporal dependencies across trials, a temporal generalization matrix is constructed based on the processed data from Section 2.2, using a time window size of 100 ms and a 25% overlap, following the approach of Veillette et al. ^(26)^. To classify frequent versus infrequent stimuli in this matrix, a model was trained using a sliding window of three consecutive time points (i.e., a target time point and its immediate neighbors), and predictions were made by shifting the test time point. Classification performance was assessed using the F1 score (Equation 1). This matrix aims to identify the most suitable time periods for classification. While Veillette et al. ^(26)^ used logistic regression to classify EEG signals related to the sense of agency during self-executed and unexecuted motor tasks based on electromyogram information, this study employed a Support Vector Machine (SVM), which also allows for nonlinear classification, to evaluate the performance. The F1 score represents the balance between precision and recall that increases with a higher number of true positives (TP) and decreases when false positives (FP) or false negatives (FN) increase. TP refers to correctly classified frequent stimuli, FP to infrequent stimuli misclassified as frequent, TN to correctly classified infrequent stimuli, and FN to frequently misclassified stimuli.

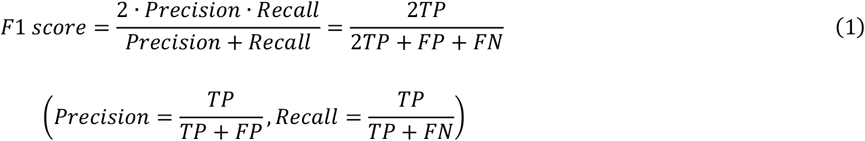

#### 2.4.2. Classification using six models

Based on the results in Section 2.4.1, the analysis was restricted to periods suitable for classification, and six different models were trained to predict whether each trial corresponded to a frequent or infrequent stimulus. The F1 score is used as the evaluation metric. For this analysis, the time window size was set to 10 ms with a 10% overlap. Although this differs from the previous window size, a smaller window is expected to yield a higher precision. The six classification models used were SVM, Random Forest, Logistic Regression, XGBoost, CatBoost, and K-means. These models were selected based on previous studies ^(19)∼(21)^, emphasizing those known for their high classification performance and computational efficiency. The model performance was evaluated using cross-validation. For each fold, the training data were further split into training and validation sets using 5-fold cross-validation to optimize the hyperparameters. The classifier was then retrained on the full training set using the best parameters, and predictions were made on the test set, which comprised 20% of the total data. The F1 scores of the five folds were averaged to obtain the final mean F1 score.

#### 2.4.3. Evaluation

The performances of the classification models were evaluated using the area under the receiver operating characteristic (ROC) curve (AUC). According to Hanley and McNeil ^(27)^, the AUC represents the probability that a randomly selected positive instance is ranked higher than a randomly selected negative instance, with 0.5 indicating chance-level performance. To assess the effects of the three experimental factors described in Section 1.3, namely the modality of infrequent stimuli, a combination of frequent and infrequent stimuli, and the response type to infrequent stimuli, a three-way mixed-design analysis of variance (ANOVA) was conducted. This analysis used the average F1 score of the classification model that achieved the most stable and accurate AUC performance. To evaluate the influence of polarity on classification outcomes, we also calculated the F1 scores using microstate templates that did not account for polarity. Furthermore, to examine whether the classification performance depended on the number of templates, we computed the F1 scores under three additional conditions: (1) using four clusters, (2) using seven clusters derived in a data-driven manner (7_dat), and (3) using five clusters with two additional artificially created templates generated by averaging the centroids of microstates A and B, and microstates D and E, respectively (7_art). These results were analyzed using ANOVA.

## 3. Results

### 3.1. Label Occurrence Frequency

Fig. 2 shows the average label occurrence rates for EEG microstate labels across participants in the uniV-only condition. In Fig. 2(a), during the keyresTask, infrequent visual stimuli elicited higher occurrence rates of C+ and E+ than frequent stimuli within the first 200 ms following stimulus onset. This was followed by a transition to D+, C−, and E− around 300–400 ms, and a return to C+ and E+ after 500 ms. A similar pattern was observed in the countTask (Fig. 2(b)).

**Fig. 2.**
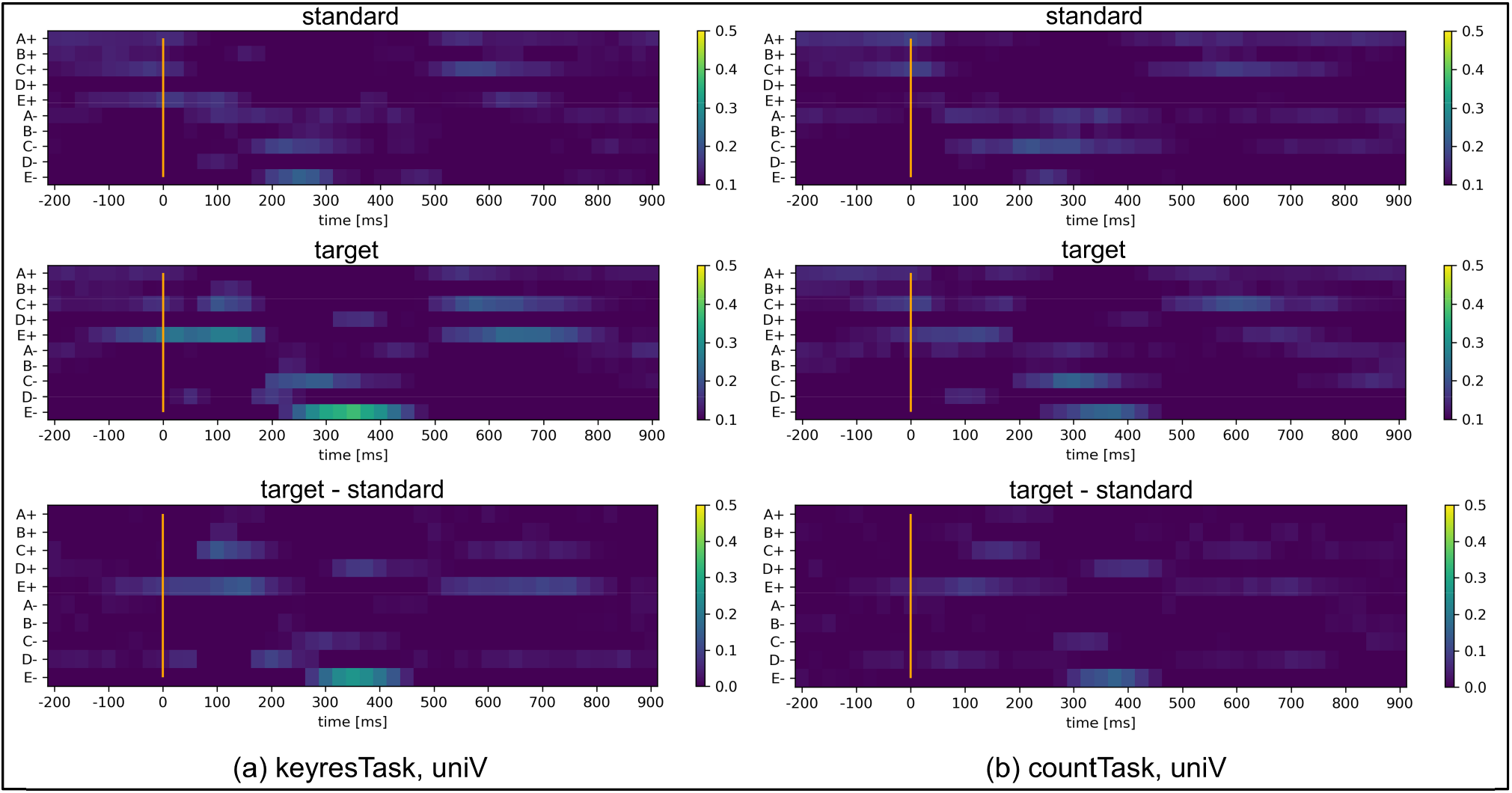
EEG microstate occurrence rates for the uniV condition in the keyresTask and countTask. (a) Results for the key response task. From top to bottom: the occurrence rate during. Standard stimuli, the occurrence rate during target stimuli, and the difference between target and standard stimuli. The horizontal axis represents time (ms), the vertical axis shows the template labels, and the color bar indicates the occurrence rate, with values shifting from dark blue to yellow as the rate increases. (b) Results for the count task are shown in the same format as in (a).

### 3.2. Result of Temporal Generalization Matrix

The results of the temporal generalization matrix are shown in Fig. 3. As seen in Fig. 3(a) and (c), F1 scores were low during the pre-stimulus period (−200 to 0 ms), and showed a clear diagonal pattern, with higher scores when the training and testing time points corresponded. Notably, training on data from the 0 to 250 ms range increased F1 scores when testing on data after 500 ms. Conversely, training on data after 500 ms improved the classification for testing in the 0–250 ms range. Fig. 3(b) and (d) show that, regardless of response type, the croV condition yielded the highest F1 scores (keyresTask: F1 score at 275 ms = 0.731 ± 0.118; countTask: F1 score at 250 ms = 0.647 ± 0.075). In contrast, the uniA condition produced the lowest scores (keyresTask: F1 score at 175 ms = 0.632 ± 0.089; countTask: F1 score at 325 ms = 0.552 ± 0.074). Across all experiments, the average F1 scores across participants peaked in the 0–400 ms range. Therefore, the 0-400ms time window was used in subsequent classification analyses.

**Fig. 3.**
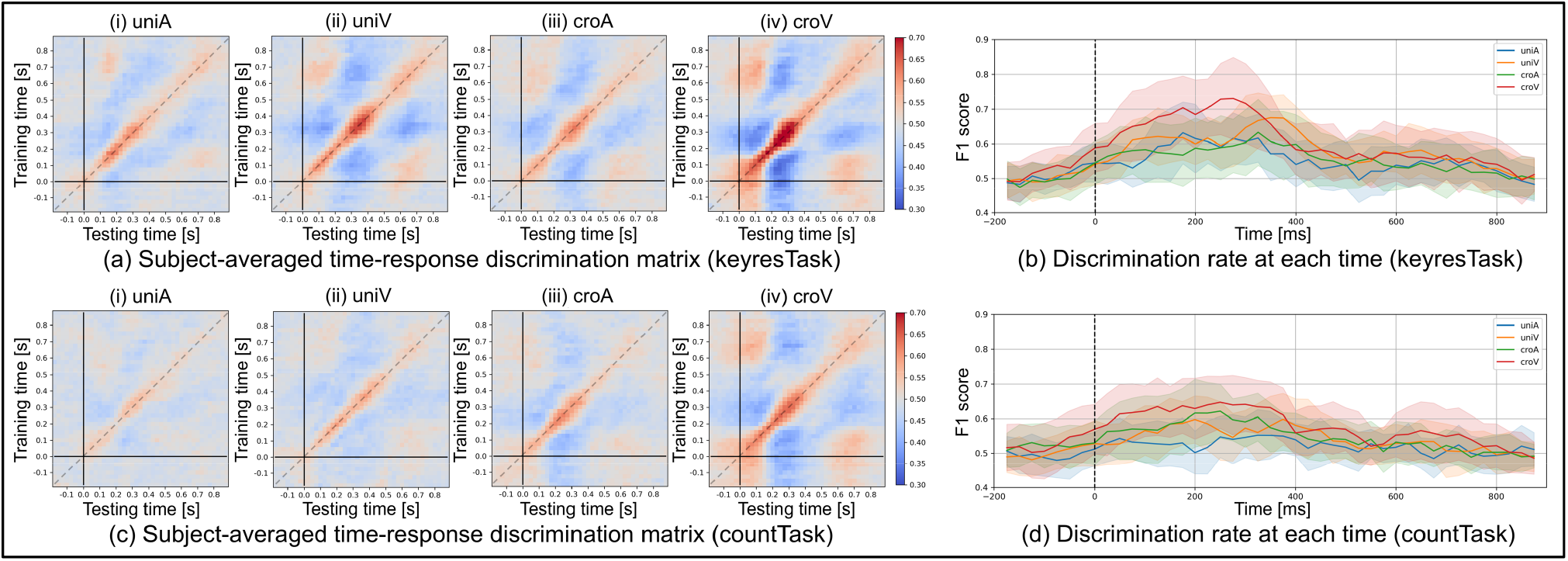
Temporal generalization matrix and discrimination rate at each time. (a) and (c) show the temporal generalization matrices averaged across all participants (n = 20) for the keyresTask and countTask, respectively (horizontal axis: testing time; vertical axis: training time). (b) and (d) display the average and standard deviation of F1 scores across participants along the diagonal of the matrices shown in (a) and (c), respectively.

### 3.3. Result of Classification Using Six Models

#### 3.3.1. Effects of stimulus modality, modality conditions, and response type

Based on the results in Section 2.4.2, six classification models were applied to the 0–400 ms time window, which showed higher F1 scores in Section 3.2. The results are shown in Fig. 4. All models, except K-means, achieved classification performance above the chance level regardless of the response type, with the highest performance observed in the croV condition. However, the F1 scores for the SVM and Logistic Regression models were close to the chance of the uniA condition in countTask.

**Fig. 4.**
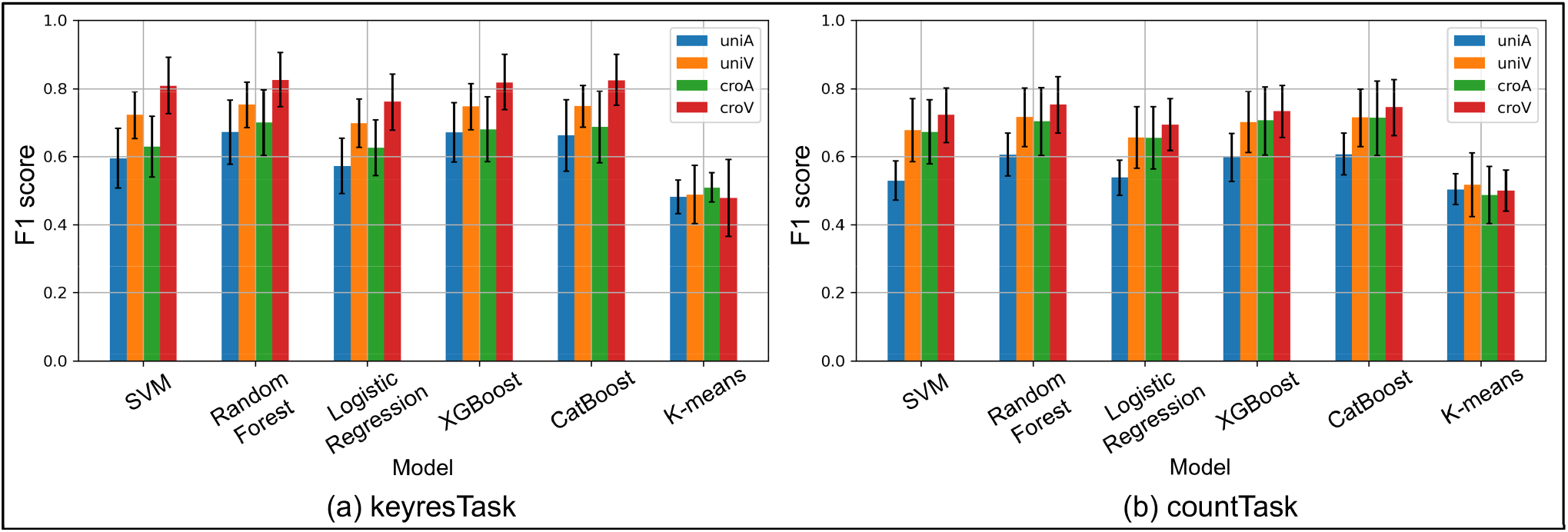
F1 scores for each machine learning model by keyresTask (a) and countTask (b). Color represents each condition (blue: uniA, orange: uniV, green: croA, red: croV). Error bar indicates standard deviation across participants.

Similar trends were observed across five classification models, excluding K-means, for both task types. A correspondence matrix was constructed to visualize these relationships (Fig. 5). Additionally, the AUC was calculated as a performance metric for each classification model. The AUC distributions for the highest-performing (croV) and lowest-performing conditions (uniA) are shown in Fig. 6, Random Forest, XGBoost, and CatBoost demonstrated the most stable and accurate classification performance. Notably, in keyresTask under the croV condition, the average AUC exceeded 0.8, even when SVM and Logistic Regression were included, indicating highly reliable classification results.

**Fig. 5.**
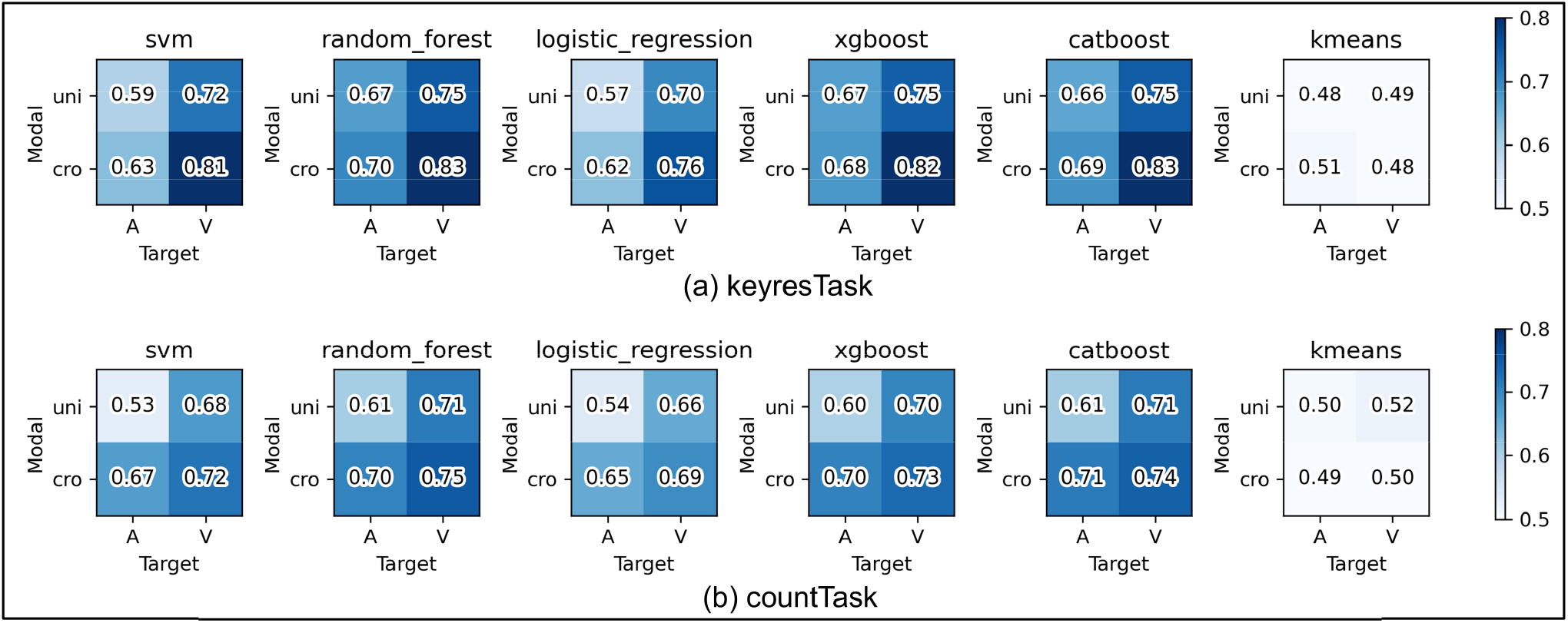
Target modal matrix by keyresTask (a) and countTask (b). The color bar indicated F1 score.

**Fig. 6.**
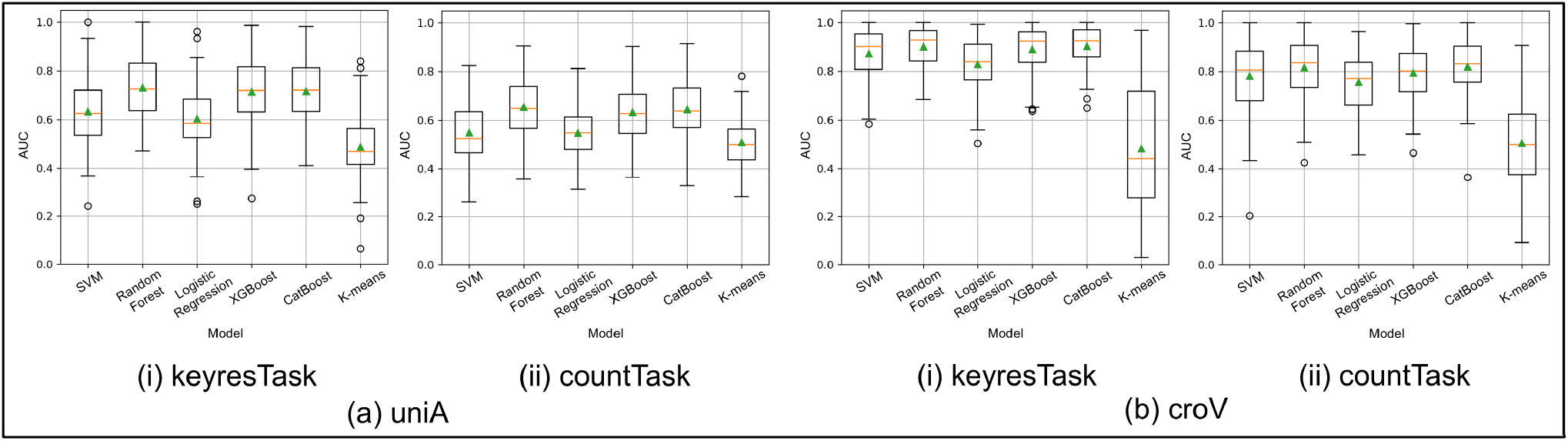
AUC box plot for each machine learning model by the lowest-performing condition (uniA (a): keyresTask (i), countTask (ii)) and the highest-performing condition (croV (b)). The vertical axis shows the AUC value.

Based on the AUC values, a three-way mixed-design ANOVA was conducted using the average scores of the three-decision tree-based models (Random Forest, XGBoost, and CatBoost), which showed the most stable and accurate performance. The analysis revealed a significant second-order interaction effect (*F*(1, 38) = 8.65, *p*<.01,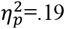). Regarding the modality of infrequent stimuli, visual stimuli yielded significantly higher scores than auditory stimuli in all conditions except for the cross-modal condition in countTask (Fig. 7(a)). For stimulus modality combinations, cross-modal conditions with infrequent visual stimuli in keyresTask showed significantly higher scores (Fig. 7(b)). Regarding the response type, keyresTask outperformed countTask under the uniA and croV conditions, indicating different effects depending on the combination (Fig. 7(c)). These results suggest that the highest classification performance was achieved under high-frequency auditory and low-frequency visual stimuli. Moreover, keyresTask enabled a more accurate classification than countTask.

**Fig. 7.**
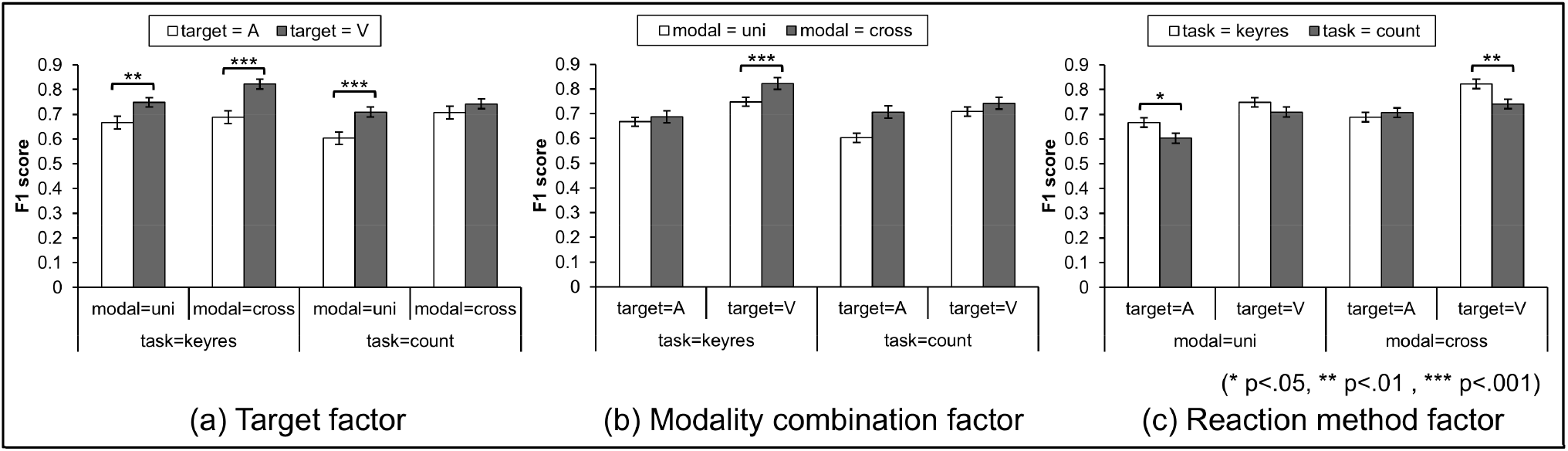
F1 scores for three experimental factors in oddball task. (a) show Target factor, (b) show Modality combination factor, and (c) show reaction method factor. The error bar represents standard deviation across participants.

#### 3.2.2. Effects of Polarity and Number of Templates

Fig. 8 presents a matrix of participant-averaged F1 scores across conditions with and without polarity and across different numbers of templates. All five models excluding K-means yielded higher classification scores when polarity-considered templates were used. Additionally, no notable changes were observed in the classification performance when the number of templates was increased or decreased.

**Fig. 8.**
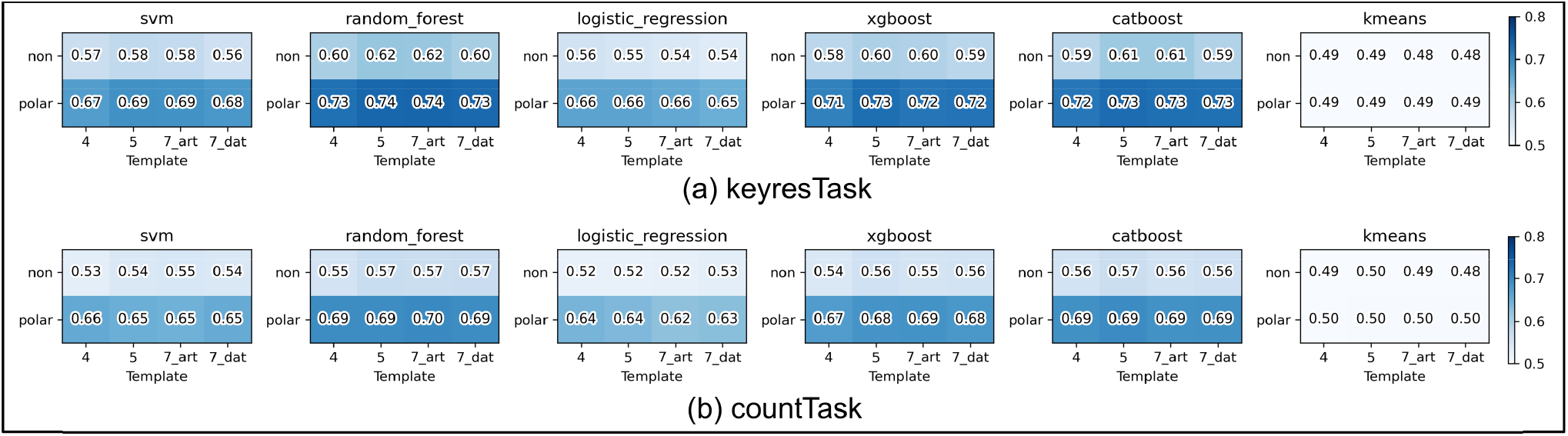
Effects of number of templates and Polarity for keyresTask (a) and countTask (b). The color bar indicated F1 score.

An ANOVA was conducted using the average F1 scores from the five classification models (excluding K-means) to examine the effects of polarity and number of template clusters, limited to cluster sizes of 4 and 5 (Fig. 9). The interaction between cluster number and polarity was not significant for either task (keyresTask: *F*(1, 19) = 0.00, *p*<.99,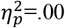; countTask: *F*(1, 19) = 3.16, *p*<.09, 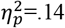). However, adding polarity significantly improved classification accuracy in both tasks (keyresTask: 20.1% improvement, *F*(1, 19) = 182.96, *p*<.001, 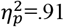; countTask: 22.2% improvement, *F*(1, 19) = 368.96, *p*<.001, 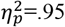). Regarding the number of clusters, the keyresTask showed significantly higher performance with five clusters compared to four (*F*(1, 19) = 9.32, *p*<.01, 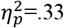).

**Fig. 9.**
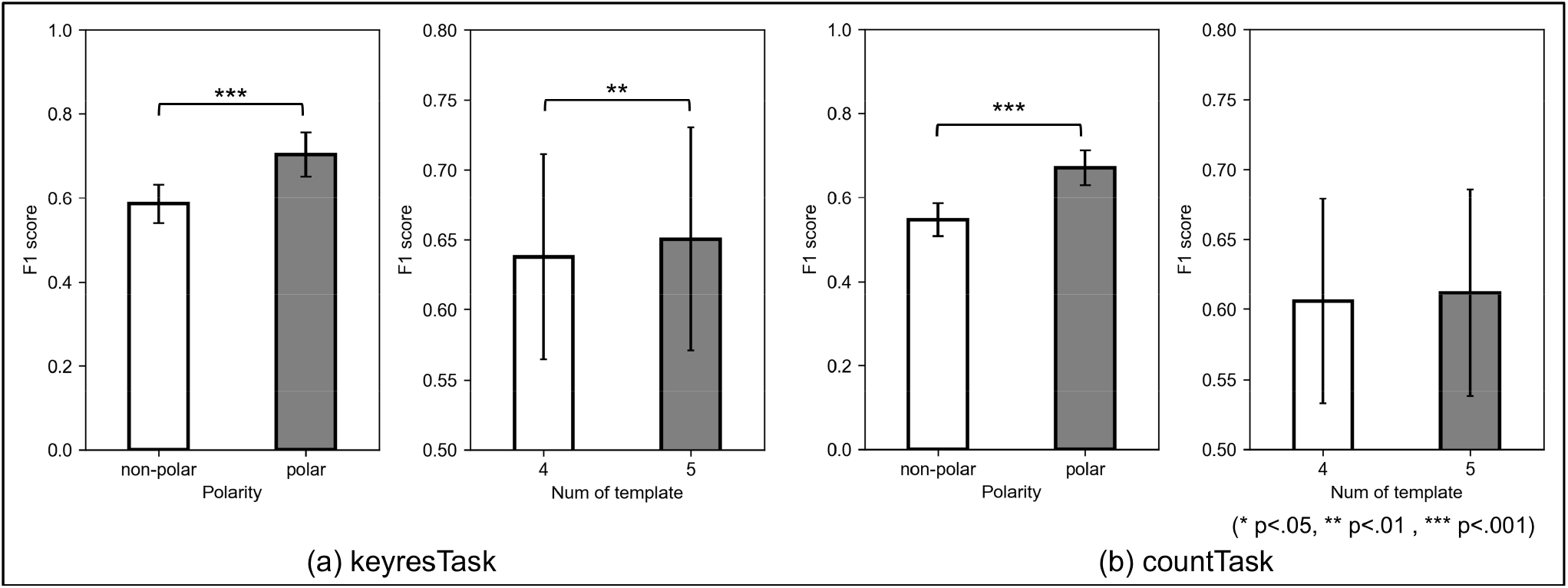
F1 scores of polarity and number of templates for keyresTask (a) and countTask (b). The error bar represents standard deviation across participants.

## 4. Discussion

This study had three main objectives: (I) evaluate the classification performance of P300 responses using microstate labeling, which differs from conventional methods; (II) explore the potential improvement in performance using polarity-considered microstate labeling; and (III) examine the relationship between classification performance and different experimental conditions in an oddball paradigm using microstate labeling. The discussion is structured around these objectives.

### 4.1 P300 Classification Performance Using a Non-Conventional Labeling Method

Unlike conventional methods such as SWLDA, which have demonstrated over 90% classification accuracy (Krusienski et al.) ^(3)^, the present study employed one-dimensional microstate label sequences for classification. Among the five models, SVM, Random Forest, Logistic Regression, XGBoost, and CatBoost achieved F1 scores above the chance level. Logistic Regression and SVM with a linear kernel can be regarded as linear classifiers. Tree-based models such as Random Forest, XGBoost, and CatBoost, which are nonlinear classifiers, showed consistently high and stable AUC values, suggesting that these models are best suited for classification based on EEG microstate labeling. The superior performance of nonlinear models compared with linear models implies that the distribution of microstate labels is inherently nonlinear.

Fig. 2 shows that labels C and E frequently appeared after stimulus onset across all tasks, but no single label dominated, indicating variability across conditions. Tarailis et al. ^(28)^, associated microstate C with internal processes such as self-reflection and autobiographical memory, whereas microstate E is linked to emotional processing and interoceptive attention. The co-occurrence of both labels suggests that the oddball task evokes parallel internal cognitive and bodily responses, resulting in dynamic transitions in brain states and a nonlinear structure in the label sequences. Even in frequent stimulus trials, label sequences occasionally resembled those of infrequent stimuli, further supporting the nonlinearity of the label distribution. Although the classification accuracy of this method reached approximately 80% at best under the croV condition, which was lower than that reported for conventional methods, the use of label sequences enabled greater interpretability of spatial dynamics in EEG, as discussed in Section 3.1.

### 4.2. Potential Performance Improvement Through Polarity-considered Microstate Labeling

This study adopted a labeling method that preserved the polarity of EEG microstates. Incorporating polarity into the templates resulted in statistically significant improvements in the classification accuracy compared to the polarity-agnostic condition. To explore how the inclusion of polarity affects label information, Fig. 10 presents the time course of the occurrence rates averaged across participants for keyresTask, following the same approach as in Section 2.3. Fig. 10 illustrates that the polarity-considered labeling produced a label space with approximately twice the granularity compared to polarity-agnostic labeling, effectively distributing the information across a broader scale and potentially enhancing the classification performance. Additionally, the underlying neural dynamics can be interpreted as follows: following the onset of an infrequent stimulus, positive frontal topographies (C+ and E+) emerged within 0–200 ms, followed by D− at 200–300 ms, and E− at 300–400 ms. Among these, D− in the N2 time window and E− in the P3 time window exhibit spatial topographies similar to the N2 and P3 components reported by Mahini et al. ^(9)^. Mahini et al. ^(9)^ stated that “Cluster 2, which represents the P3 component, was most frequently observed between 280 and 540 ms,” indicating the time window with the highest spatial correlation for the P3 cluster derived via clustering. Although the results were averaged across the participants and covered a slightly narrower time window, they exhibited comparable temporal patterns. The dynamic changes observed include initial attentional switching and early sensory response within 0–200 ms, internal re-evaluation around 200–300 ms, and evaluative or decision-related processes between 300 and 400 ms, which support the interpretation that P300 is closely linked to the E− label. However, when the polarity is ignored, this nuanced transition is lost, and these distinct states collapse into a single E label, obscuring their interpretability. This likely contributed to the observed drop in classification performance under polarity-agnostic conditions. Mahini et al. ^(9)^ also observed that polarity-inverted topographies enable the formation of stable clusters for N2 and P3, supporting the notion that polarity enhances classification precision. Similarly, as Kashihara et al. ^(8)^ pointed out, including polarity is essential for accurately capturing the continuous topological structure of the EEG state space and maintaining the inherently smooth and directional trajectories of microstate transitions. Our findings align with this perspective, suggesting that polarity information is crucial in discretely approximating nonlinear and continuous brain activity dynamics.

**Fig. 10.**
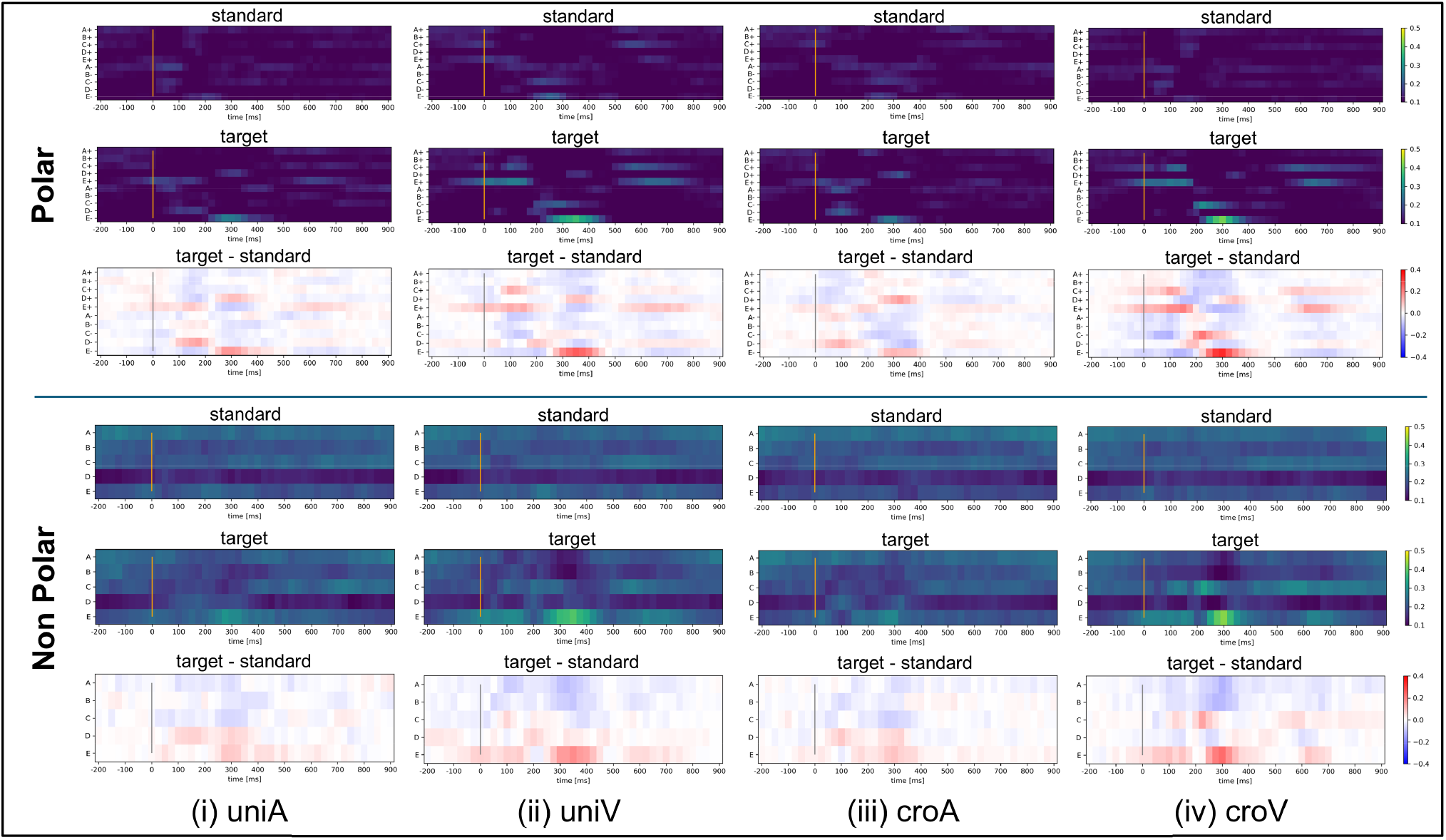
Polarity differences in EEG microstate appearance rate by keyresTask.

### 4.3 Correspondence Between Classification Performance and Oddball Task Conditions via Microstate Labeling

The effects of the three experimental factors described in Section 1.3 were evaluated and are presented in the results of Section 3.3.1.

First, regarding factor (i), stimulus modality—visual stimuli showed significantly higher classification accuracy than auditory stimuli (except in the cross-modal condition of countTask). This suggests that the visual stimuli are more effectively classified using the proposed method. These results are consistent with the findings of Katayama and Polich ^(12)^ and Polich and Heine ^(13)^. Therefore, this result may reflect occipital neural activity induced by visual input, which strongly contributes to spatial distribution patterns and forms brain states that are easier for the classifier to distinguish.

Second, for factor (ii), the combination of stimulus modalities—the cross-modal condition involving infrequent visual stimuli in the keyresTask–yielded significantly higher accuracy than the unimodal condition, while no significant differences were observed in other tasks. This indicates that the combination of modalities is effective only when the task involves rapid motor responses, such as in the keyresTask. In this study, several results aligned with findings from the traditional P300 literature. The improved classification accuracy observed in some cross-modal conditions compared with unimodal conditions may be explained by Calvert et al. ^(29)^, who reported that multimodal integration imposes an additional neural processing load. The keyresTask, which requires quick responses, may have been more influenced by this transient high cognitive load than the countTask, in which no reaction time constraint was imposed.

Finally, for factor (iii), response type to infrequent stimuli, a significantly higher classification accuracy was observed in the keyresTask than in the countTask in the uniA (auditory-only) and croV (frequent auditory, infrequent visual) conditions. This indicates that the task difference emerged only when the frequent stimulus was auditory. These findings are consistent with those reported by Brázdil et al. ^(17)^ and Kotchoubey ^(18)^. One likely explanation for this task-related difference is that keyresTask requires participants to respond quickly, whereas countTask does not, resulting in varied response times across trials in the latter. Additionally, the task differences appearing only when the frequent stimulus was auditory may be because auditory stimuli tend to elicit faster responses than visual stimuli. In a study by Shelton and Kumar ^(30)^, the mean reaction time for auditory stimuli was reported to be approximately 284 ms, significantly shorter than the 331 ms observed for visual stimuli. This finding supports the interpretation that keyresTask, which involves immediate responses, yields a significantly higher classification accuracy under auditory-dominant conditions.

### 4.4 Limitations and Future Directions

Despite the promising results, this study has several limitations. First, since the microstate templates used in this study were derived from the LEMON dataset ^(25)^, individual variability in brain state dynamics may have been overlooked. Utilizing subject-specific templates could further enhance classification accuracy. Second, because this study focused on offline analysis using multiple classification models, the classification performance in real-time BCI systems remains unverified. Future studies should explore the practical implementation of the proposed method to real-time BCI systems, such as the P300 speller.

## 5. Conclusion

This study examined the classification accuracy of infrequent stimuli in an oddball paradigm using polarity-considered microstate labeling and explored its applicability to BCI systems. Classification models, excluding K-means, achieved significantly higher accuracy when polarity was incorporated into the labeling process. However, the number of templates had a limited effect on performance. Notably, the cross-modal condition with infrequent visual stimuli in keyresTask yielded the highest classification accuracy, suggesting that both the response type and stimulus modality influence the effectiveness of microstate-based classification. These findings indicate that using one-dimensional polarity-considered microstate labeling for EEG signals is a promising approach for BCI applications, including P300 spellers. Moreover, performance may be further enhanced by optimizing experimental conditions such as stimulus modality and response type.

## Acknowledgements

We would like to thank Editage (www.editage.jp) for English language editing.

